# Monitoring ligand-induced changes in receptor conformation with NanoBiT conjugated nanobodies

**DOI:** 10.1101/2020.04.21.032086

**Authors:** Mark Soave, Raimond Heukers, Barrie Kellam, Jeanette Woolard, Martine J. Smit, Stephen J. Briddon, Stephen J. Hill

**Affiliations:** Division of Physiology, Pharmacology and Neuroscience, School of Life Sciences, University of Nottingham, Nottingham, NG7 2UH, UK; Centre of Membrane Proteins and Receptors (COMPARE), University of Birmingham and University of Nottingham, The Midlands, UK; Division of Medicinal Chemistry, Amsterdam Institute for Molecules, Medicines and Systems (AIMMS), VU University of Amsterdam, De Boelelaan 1108, 1081 HZ Amsterdam, The Netherlands; QVQ Holding B.V., Yalelaan 1, 3584 CL Utrecht, The Netherlands; School of Pharmacy, Centre for Biomolecular Sciences, University of Nottingham, Nottingham, NG7 2RD, UK

**Keywords:** Nanobodies, chemokine receptor, CXCR4, NanoBiT, NanoLuciferase, conformational selectivity, extracellular loop 2

## Abstract

Camelid single-domain antibody fragments (nanobodies) offer the specificity of an antibody in a single 15kDa immunoglobulin domain. Their small size allows for easy genetic manipulation of the nanobody sequence to incorporate protein tags, facilitating their use as biochemical probes. The nanobody VUN400, which recognises the second extracellular loop of the human CXCR4 chemokine receptor, was used as a probe to monitor specific CXCR4 conformations. VUN400 was fused via its C-terminus to the 11-amino acid HiBiT tag (VUN400-HiBiT) which complements to LgBiT protein, forming a full length functional NanoLuc luciferase. Here, complemented luminescence was used to detect VUN400-HiBiT binding to CXCR4 receptors expressed in living HEK293 cells. VUN400-HiBiT binding to CXCR4 could be prevented by orthosteric and allosteric ligands, allowing VUN400-HiBiT to be used as a probe to detect specific conformations of CXCR4. These data demonstrate that the high specificity offered by extracellular-targeted nanobodies can be utilised to probe receptor pharmacology.

## Introduction

The C-X-C chemokine receptor 4 (CXCR4) is a family A G protein-coupled receptor (GPCR) which plays an important role in the immune response and in the progression of many diseases, including cancer and HIV infection (Kucia et al., 2004; Scholten et al., 2012). As a GPCR, CXCR4 consists of seven transmembrane helices, an extracellular N-terminus and an intracellular C-terminus. Of the chemokine receptor family, CXCR4 is unusual in that it exclusively binds one chemokine ligand, CXCL12 (formerly termed stromal cell-derived factor 1α, SDF-1α). CXCL12 binds to CXCR4 in a two-step process: first interacting with residues in the N-terminus of the receptor (chemokine recognition site 1, CR1), before CXCL12 engages both the extracellular loops and binding pocket within the transmembrane helices (chemokine recognition site 2, CR2; Kofuku et al., 2009). Mutation studies have shown that the N-terminus of the receptor contributes to chemokine binding, whereas the binding pocket within the transmembrane helices is responsible for receptor activation and binding affinity (Scholten et al., 2012). The structure of CXCR4 has since been solved in the presence of the small molecule antagonist IT1t, the cyclic peptide antagonist CVX15 (Wu et al., 2010), and the chemokine antagonist vMIP-II (Qin et al., 2015). Furthermore, CXCR4-targeting small molecules bind to a site which partially or wholly overlaps with CR2 (Arimont et al., 2017). Although CXCR4 only binds one type of chemokine, CXCL12, these studies have shown that CXCR4 is able to adopt many different conformations in order to accommodate and bind these different classes of ligands.

Due to the critical role CXCR4 plays in HIV infection and tumour progression, there has been considerable interest in developing specific CXCR4 antagonists, a number of which are in the clinical development pipeline (Kuhne et al., 2013; www.clinicaltrials.gov). Despite this, only the small molecule CXCR4 antagonist AMD3100 (Plerixafor, Mozobil®) has received FDA approval for the treatment of lymphoma and multiple myeloma (Keating, 2011), and the treatment of Hodgkin lymphoma in Europe (Douglas et al., 2018). AMD3100 has also been demonstrated to be a weak allosteric agonist of the atypical chemokine receptor 3 (ACKR3; previously CXCR7), inducing β-arrestin recruitment *in vitro* (Kalatskaya et al., 2009). This, in addition to its short serum half-life *in vivo* (Hendrix et al., 2000), has necessitated the development of more selective, long-lasting antagonists. The improved selectivity and extended half-lives of antibodies compared to small molecules has meant there has been much interest in using antibody-based approaches to target CXCR4 therapeutically (Hutchings et al., 2017; Bobkov et al., 2019). This has included the recent development of a panel of single domain antibody fragments, called nanobodies, which are able to bind CXCR4 (Jahnichen et al., 2010; de Wit et al., 2017; Bobkov et al., 2018; Van Hout et al., 2018).

Nanobodies are small proteins (circa 12-15 kDa), derived from the single variable fragments (V_HH_) of heavy-chain only antibodies found in members of the Camelidae family. Nanobodies are known to be excellent conformational sensors due to their small size and three-dimensional structure (De Genst et al., 2006). Furthermore, their elongated complimentary determining region 3 (CDR3) enables nanobodies to engage hidden cavities and conformational epitopes (De Genst et al., 2006). Nanobodies have been extensively used within the GPCR field to stabilise specific receptor conformations for crystallization (Rasmussen et al., 2011; Ring et al., 2013; Kruse et al., 2013; Huang et al., 2015; Che et al., 2018), and to elucidate new conformational states (Staus et al., 2016). This has also led to the development of these nanobodies as biosensors to investigate GPCR signalling (Irannejad et al., 2013; Staus et al., 2014; Staus et al., 2016; Stoeber et al. 2018). The nanobodies used in these studies generally target intracellular regions of the GPCR, often binding in the same pocket as G proteins to act as G protein mimetics (Rasmussen et al., 2011; Staus et al., 2014; Stoeber et al., 2018).

Nanobodies which that bind to the extracellular domains of GPCRs are able to modulate receptor activity and organization (De Groof et al., 2019a). Several studies have investigated the therapeutic potential of extracellular nanobodies which target chemokine GPCRs, including CXCR2 (Bradley et al., 2015), CXCR4 (Jahnichen et al., 2010; de Wit et al., 2017; Bobkov et al., 2018; Van Hout et al., 2018), ACKR3 (Maussang et al., 2013), and US28 (De Groof et al., 2019b). Given their relatively large N-terminus compared to the other Class A GPCRs, and the fact that their endogenous ligands are peptides, chemokine GPCRs are ideal candidates to target with extracellular nanobodies. Most recently, several nanobodies binding to the N-terminus and second extracellular loop (ECL2) of CXCR4 were generated (Bobkov et al., 2018; Van Hout et al., 2018). For example, VUN400 was one of these nanobodies that acted as an antagonist and inhibited CXCL12-induced signalling by CXCR4, as well as internalization. Interestingly, VUN400 also showed a decreased potency of inhibiting CXCR4-medicated HIV-1 entry compared to its ability to inhibit CXCL12-induced signalling, suggesting a conformational sensitivity of the nanobody (Van Hout et al., 2018).

The recently developed NanoLuc binary technology (NanoBiT) splits the bright NanoLuc luciferase into two segments at the C-terminal region, the 18 kDa fragment (termed LgBiT), and the 1.3 kDa small complementation tag (termed SmBiT; Dixon et al., 2016). These fragments have low intrinsic affinity and complement to form the full luminescent NanoLuc protein but with a reduced luminescence compared to the full length NanoLuc (Dixon et al., 2016). Other small complementary peptides with a range of affinities for LgBiT were identified, including an 11 amino acid sequence with very high affinity, termed HiBiT. The complemented HiBiT-LgBiT protein showed a similar luminescence output to the full length NanoLuc, making it an ideal system to study proteins expressed at endogenous levels (Schwinn et al., 2018). NanoBiT has been used to monitor protein-protein interactions, including GPCR oligomerization (Botta et al., 2019), and the recruitment of G proteins and β-arrestin to GPCRs (Hisano et al., 2018; Laschet et al., 2018; Storme et al., 2018), with the rapid complementation and maturation rate of the split NanoLuc luciferase enabling kinetic measurements. These studies made use of GPCRs with NanoBiT fused to their C-terminal domains. In addition, we have recently demonstrated the use of N-terminally-fused NanoBiT to monitor adenosine A_1_ receptor internalization *in vitro* (Soave et al., 2019b), showing it was possible to use NanoBiT to measure the membrane expression of GPCRs in living cells.

The HiBiT-LgBiT system has also recently been used to monitor conformational changes in CXCR4 following CRISPR/Cas9 mediated homology directed repair to insert HiBiT onto the N-terminus of CXCR4 within the endogenous genome (White et al., 2020). This study showed that AMD3100 induced a marked increase in luminescence following complementation of the HiBiT attached to the N-terminus of endogenous CXCR4 with purified LgBiT (White et al., 2020). This was proposed to be a consequence of changes in affinity of HiBiT for LgBiT as a result of extracellular conformational changes in CXCR4 induced by AMD3100, reducing the steric hindrance imparted by the receptor to which HiBiT was attached (White et al., 2020).

Here, we have used the NanoBiT technology in combination with the extracellular loop 2-directed nanobody VUN400 to directly monitor changes in the conformation of CXCR4 induced by AMD3100, CXCL12, and IT1t in living cells.

## Results

### Characterisation of LgBiT-CXCR4 with NanoBiT

As part of the necessary tool development, we initially used the NanoBiT technology to characterise the binding of VUN400 to CXCR4. To establish this technique, we first modified the N-terminus of the human CXCR4 receptor through the fusion of the large NanoLuc subunit (LgBiT, Dixon et al., 2016). This receptor (LgBiT-CXCR4) was successfully expressed at the plasma membrane in HEK293 cells and could be detected following the addition of 10 nM exogenous purified HiBiT by monitoring luminescence (Figure 1a-c). HiBiT was added in the form of a purified HiBiT-HaloTag fusion protein, *circa* 34 kDa in size, which was too large to cross the plasma membrane. There was significantly increased luminescence in HEK293 cells stably expressing LgBiT-CXCR4 compared to untransfected HEK293 cells (Figure 1b). The continuous monitoring of complemented luminescence allowed for the kinetic rate constants for HiBiT at the LgBiT-CXCR4 to be determined (*k*_*on*_ 6.03×10^5^ ± 0.97×10^5^ M^−1^ min^−1^; *k*_*off*_ 0.22 ± 0.03 min^−1^; pK_D_ 6.43 ± 0.13; n=3; Figure 1d). However, it was noticeable that there was a small drop in luminescence signal after about 10min with higher concentrations of purified HiBiT (Figure 1d).

**Figure 1.**
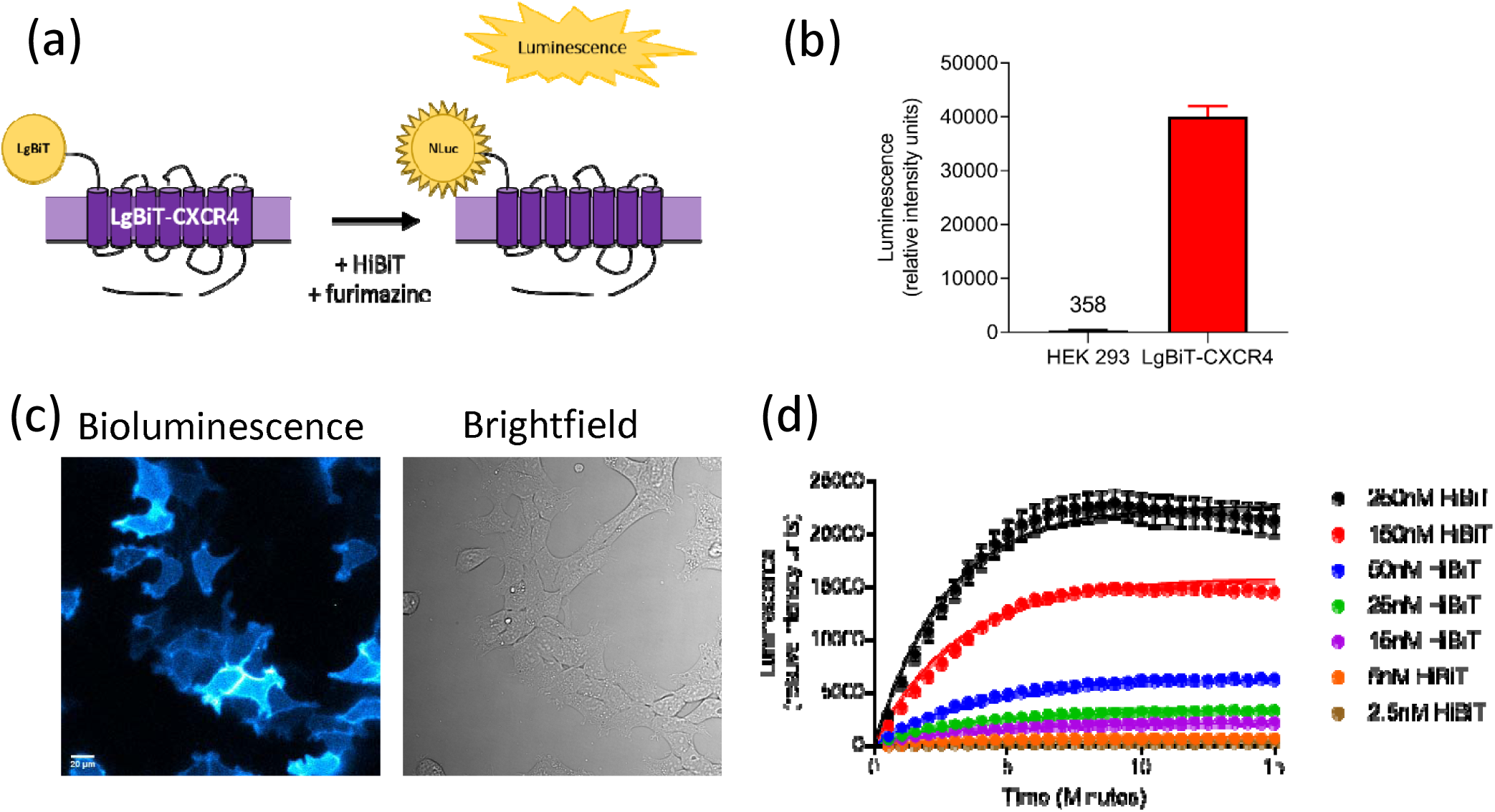
Luminescence to monitor plasma membrane expression of LgBiT-CXCR4. (a) Schematic of NanoBiT complementation to detect receptor expression. (b) PHERAstar-detected luminescence or (c) bioluminescent imaging of HEK293 cells or HEK293 cells stably expressing LgBiT-CXCR4 following treatment with 20 nM HiBiT-HaloTag and furimazine. Scale bar in (c) is 20 µm. (d) Complemented luminescence of LgBiT-CXCR4 over time following addition of multiple HiBiT concentrations. These single experiments are representative of (b) four, (c) five, or (d) three separate experiments.

The addition of purified HiBiT to LgBiT-CXCR4 resulted in the fully complemented NanoLuc-CXCR4 (Figure 2a), thus allowing NanoBRET to be performed to monitor ligand binding at the plasma membrane. Specific binding of fluorescently-labelled CXCL12 (CXCL12-AF647; Van Hout et al., 2018) could be detected at the LgBiT-CXCR4 using the previously described NanoBRET ligand binding (Stoddart et al. 2015) (pK_D_ 7.80 ± 0.08; n=4, Figure 2b). These data demonstrate that the LgBiT-CXCR4 receptor was expressed at the plasma membrane, the LgBiT-tag did not interfere with ligand binding, and the LgBiT-CXCR4 receptor was able to bind the fluorescent orthosteric agonist with high affinity.

**Figure 2.**
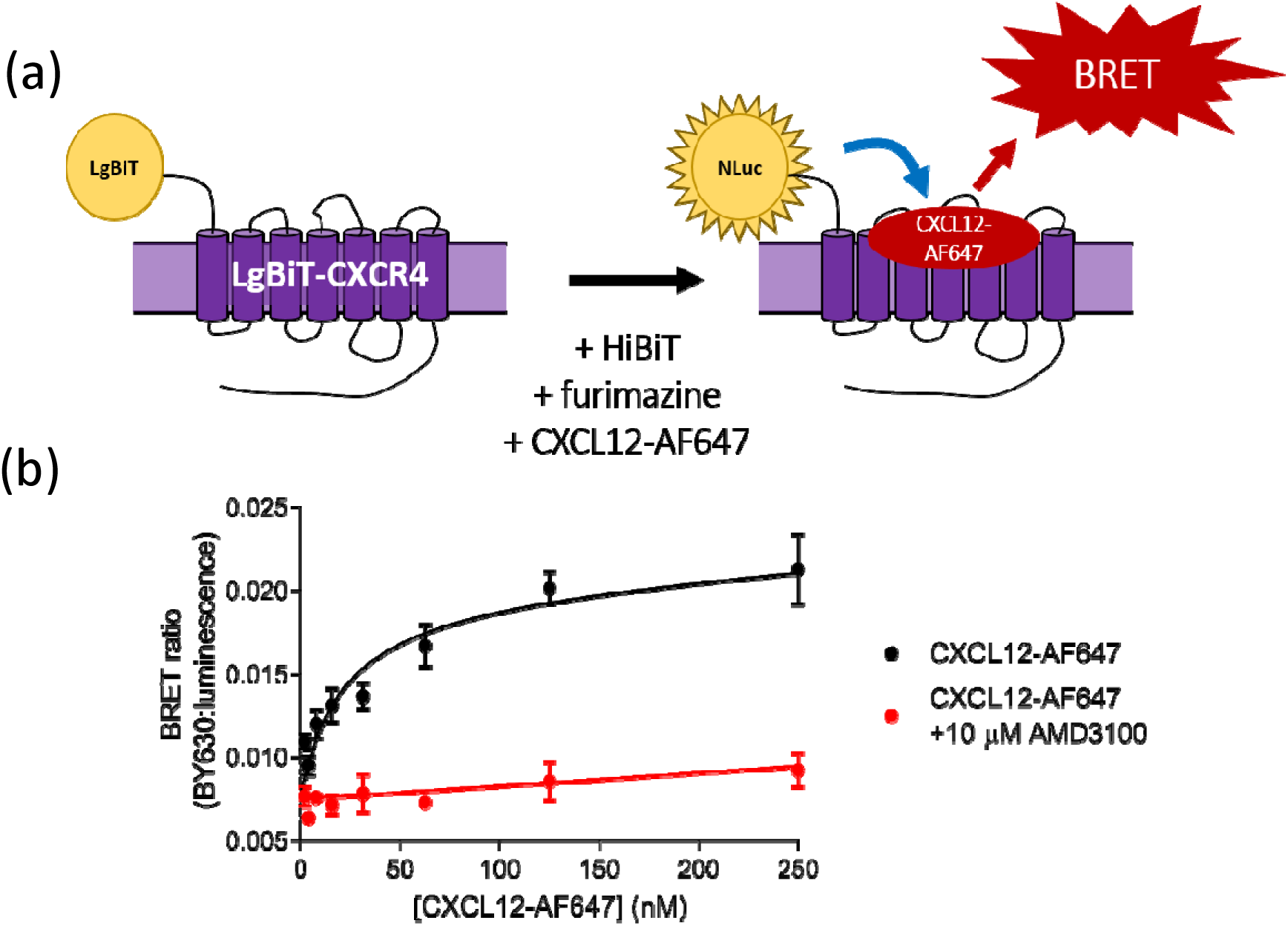
Complemented NanoBiT to perform NanoBRET ligand binding. (a) Schematic of NanoBiT complementation in order to perform NanoBRET ligand binding. (b) NanoBRET saturation binding of CXCL12-AF647 in the absence (black circles) or presence (red circles) of 10 µM AMD3100 at complemented LgBiT-CXCR4. Data are mean ± SEM from triplicate determinations in a single experiment. These single experiments are representative of four separate experiments.

### Generation of HiBiT-tagged VUN400

Next, a nanobody-HiBiT fusion construct was generated. We used the previously described nanobody, VUN400, as this is known to bind to key residues in ECL2 of the CXCR4 receptor (Bobkov et al., 2018; Van Hout et al., 2018). VUN400-HiBiT was expected to bind in close proximity to the N-terminal LgBiT on the LgBiT-CXCR4 receptor, increasing the likelihood of HiBiT-LgBiT complementation upon nanobody binding to the LgBiT-CXCR4. The plasmid DNA encoding the VUN400 nanobody was modified at the 3’ end by the incorporation of the oligonucleotide HiBiT tag (amino acid sequence: VSGWRLFKKIS) in between the C-terminal Myc-and His-tags (Figure 3a), creating a VUN400-HiBiT fusion construct.

**Figure 3.**
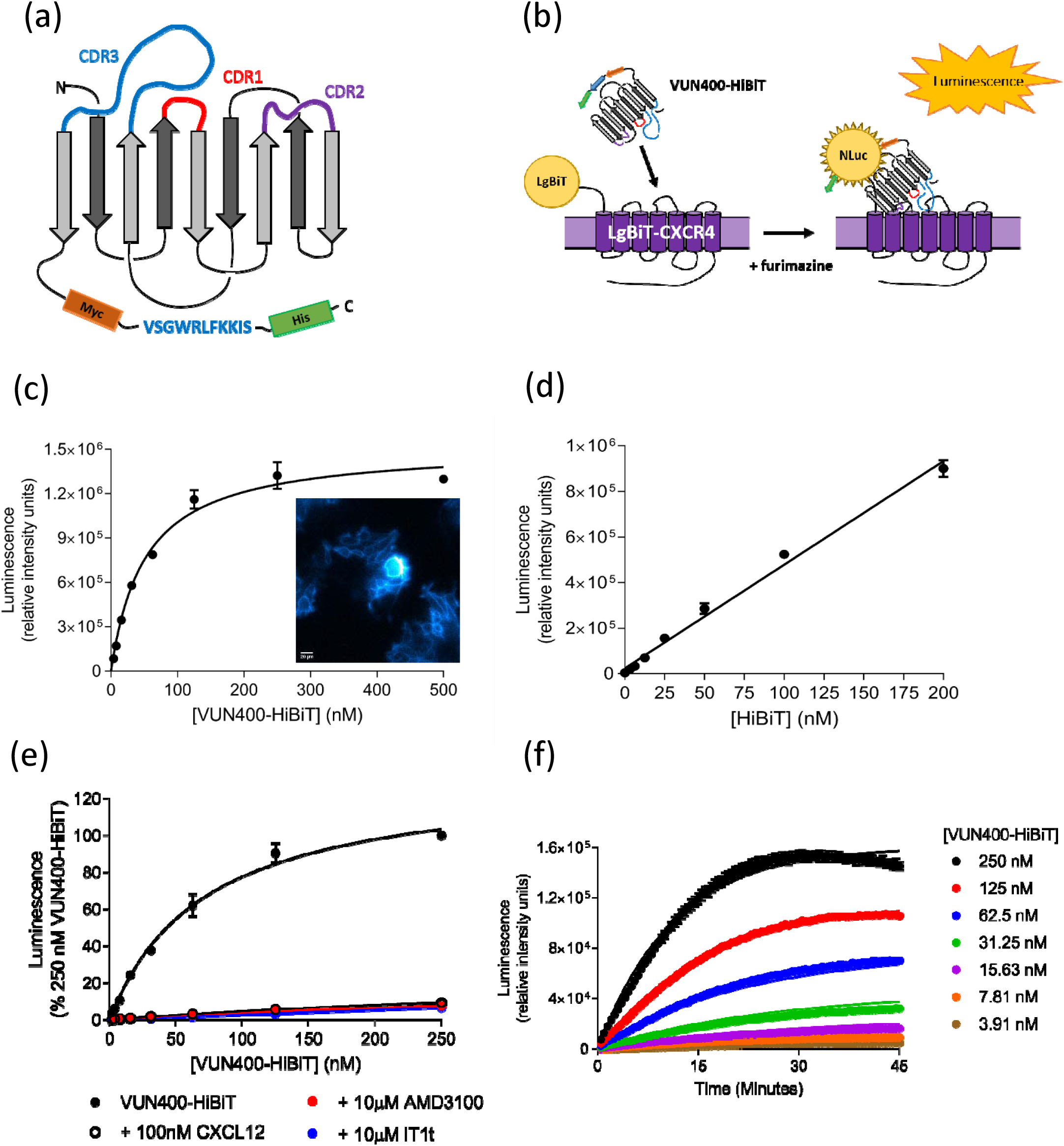
VUN400-HiBiT binding LgBiT-CXCR4. (a) Schematic of VUN400-HiBiT structure, with HiBiT amino acid tag positioned between C-proximal Myc and His tags. (b) Schematic of VUN400-HiBiT binding LgBiT-CXCR4. (c) Saturation binding of VUN400-HiBiT at the LgBiT-CXCR4. (c inset) Bioluminescent imaging of 20 nM VUN400-HiBiT binding LgBiT-CXCR4 following treatment with furimazine. (d) Saturation binding of purified HiBiT at the LgBiT-CXCR4. (e) Saturation binding of VUN400-HiBiT at LgBiT-CXCR4 in the absence (black circles) or presence of 100 nM CXCL12 (open circles), 10 µM AMD3100 (red circles), or 10 µM IT1t (blue circles). Data are expressed as a percentage of luminescence obtained with 250 nM VUN400-HiBiT alone. (f) Association binding kinetics of VUN400-HiBiT at the LgBiT-CXCR4. Data are mean ± SEM from triplicate or duplicate (+ 10 µM AMD3100 in 3e) determinations in a single experiment. These single experiments are representative of four separate experiments.

The ability for the VUN400-HiBiT to bind to the LgBiT-CXCR4 receptor was investigated thereafter (Figure 3b). Increasing concentrations of VUN400-HiBiT resulted in a saturable increase in luminescence, with an affinity of *circa* 60 nM, corresponding to a pK_D_ value of 7.25 ± 0.08, n=4 (Figure 3c). Specific binding of VUN400-HiBiT at the LgBiT-CXCR4 receptor was visualised using bioluminescence microscopy (Figure 3c inset), showing clear plasma membrane luminescence. To confirm that the observed affinity was predominantly a result of nanobody-receptor binding, HEK LgBiT-CXCR4 cells were treated with increasing concentrations of purified HiBiT (Figure 3d). Over the concentration range of purified HiBiT employed, binding was effectively linear indicative of low affinity (Figure 3d). The affinity of this interaction was significantly lower than that measured with VUN400-HiBiT, suggesting NanoBiT complementation was not driving the VUN400-HiBiT affinity measured above. VUN400-HiBiT binding to LgBiT-CXCR4 could be prevented by pre-treatment with a fixed concentration of CXCL12, as well as the antagonists AMD3100 and IT1t (Figure 3e).

Continuous monitoring of complemented luminescence made it possible to perform association kinetic experiments in order to assess the binding kinetics of VUN400-HiBiT at the LgBiT-CXCR4. As expected, complemented luminescence increased over time, and with increasing concentrations of VUN400-HiBiT (Figure 3f). Global analysis of these data produced the following kinetic rate constants (*k*_*on*_ 1.98×10^5^ ± 0.15×10^5^ M^−1^ min^−1^; *k*_*off*_ 0.032 ± 0.003 min^−1^; n=4), and the equilibrium dissociation constant determined from these rate constants (pK_D_ 6.79 ± 0.16) was in good agreement with that determined from equilibrium saturation binding experiments (Figure 3c). There was, however, a small drop in luminescence after 40 min at the highest concentrations of VUN400-HiBiT employed (Figure 3f).

### VUN400-HiBiT to detect endogenous CXCR4 receptors

The high affinity binding between VUN400-HiBiT and CXCR4 provided the opportunity to monitor ligand binding at CXCR4 receptors expressed under endogenous expression. The immortalised Jurkat T-cell line was used, as these cells express high levels of CXCR4 endogenously. Jurkat cells were treated with 100 nM VUN400-HiBiT in combination with competing ligands for two hours. Jurkat cells were then washed to remove any unbound VUN400-HiBiT, and full-length NanoLuciferase was complemented with the addition of 10 nM exogenous LgBiT. Binding of VUN400-HiBiT to endogenous, untagged CXCR4 could be detected using this setup (Figure 4a). In addition, VUN400-HiBiT could be displaced by the addition of CXCL12, AMD3100 and IT1t (Figure 4a; p<0.05), demonstrating the sensitivity of complemented luminescence as an experimental readout for monitoring nanobody-receptor binding. VUN400-HiBiT was able to probe for the differences in receptor expression between the overexpressing LgBiT-CXCR4 cell line and the Jurkat cell line. Here, the Jurkat cell line was found to express the CXCR4 receptor at lower levels than that of the LgBiT-CXCR4 cell line (15.2 ± 5.4% expression compared to LgBiT-CXCR4; n=4; Figure 4b).

**Figure 4.**
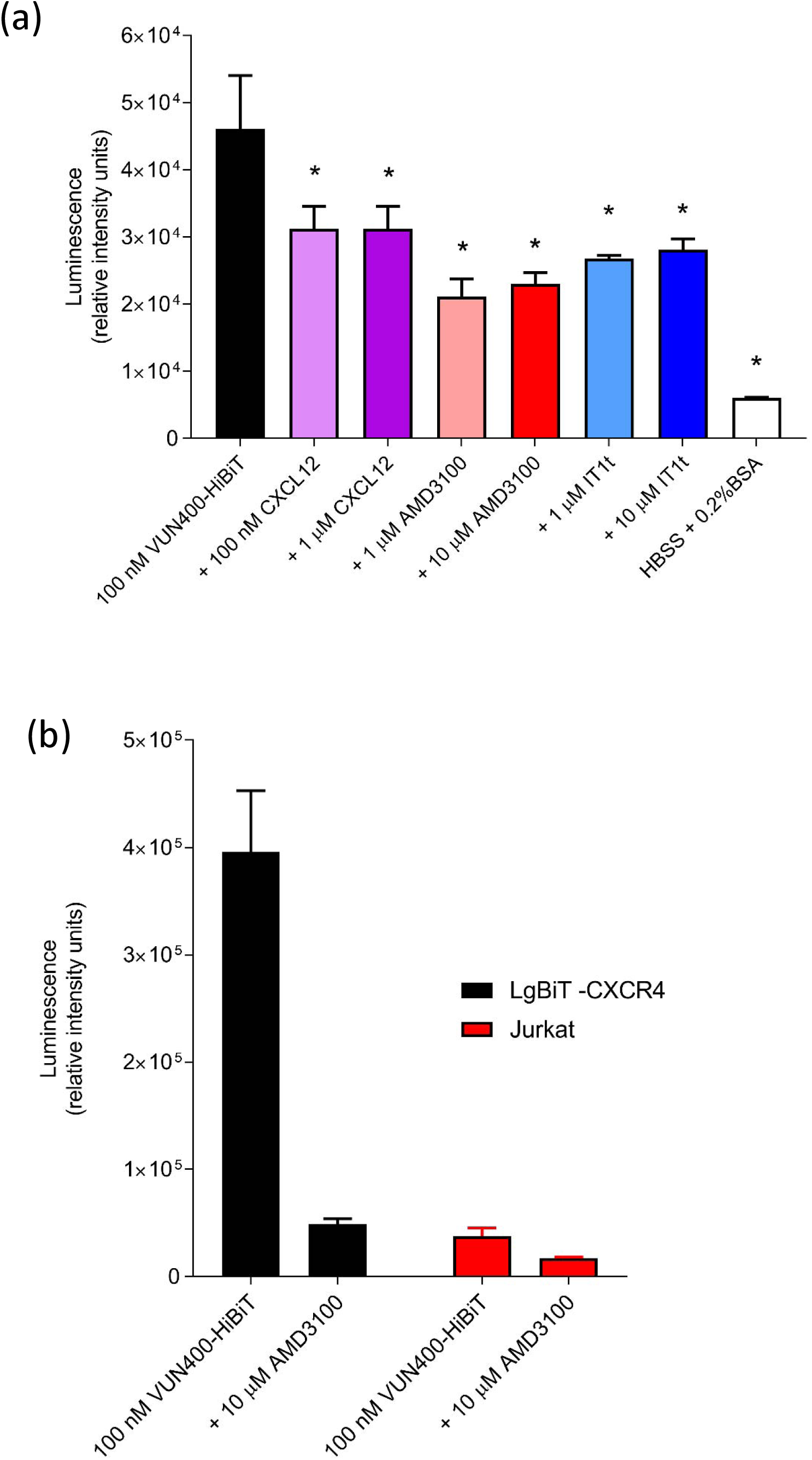
(a) VUN400-HiBiT binding in immortalised T-cells. Complemented luminescence of Jurkat cells treated with 100 nM VUN400-HiBiT in the absence or presence of CXCL12, AMD3100 or IT1t, following the addition of 20 nM purified LgBiT. Also shown is the background luminescence due to the addition of HBSS + 0.2% BSA in place of VUN400-HiBiT. Data are mean ± SEM from triplicate determinations in a single experiment. Similar data were obtained in five further experiments. * p<0.05, comparing ligand treatment with 100 nM VUN400-HiBiT treatment alone, one-way ANOVA with Dunnett’s post hoc test for multiple comparisons. Analysis of the mean data obtained in all six repeat experiments by two-way ANOVA yielded significant inhibitions (p<0.05) by all ligand concentrations apart from 100nM CXCL12. (b) Luminescence signal from LgBiT-CXCR4 or Jurkat cells treated with 100 nM VUN400-HiBiT in the absence and presence of 10 µM AMD3100. Data are mean ± SEM from triplicate determinations in a single experiment. This single experiment is representative of four separate experiments.

### VUN400-HiBiT as a probe to detect binding of CXCL12 or small molecule antagonists

The ability for ligands to interfere with VUN400-HiBiT binding to the LgBiT-CXCR4 receptor was investigated further. Whole cells were treated with 20 nM VUN400-HiBiT for 30 minutes and then challenged with a two-hour incubation with CXCR4 ligands. The orthosteric agonist CXCL12 appeared to reduce VUN400-HiBiT binding to LgBiT-CXCR4 (pIC_50_ 8.82 ± 0.12; n=5; Figure 5a; Table 1a). The antagonists AMD3100 and IT1t were also able to reduce the luminescence resulting from VUN400-HiBiT binding in a concentration-dependent manner (Figure 5a; Table 1a). This is likely to be due to a dissociation of VUN400-HiBiT from the LgBiT-CXCR4. Alternatively, the reduced signal could be a consequence of a conformational change in CXCR4 that altered the orientation of VUN400-HiBiT (binding to ECL2) with respect to LgBiT on the N-terminus of LgBiT-CXCR4, which would prevent effective complementation of the full length Nano Luciferase.

**Table 1.** Inhibition of 20 nM VUN400-HiBiT binding to whole HEK293 cells expressing either LgBiT-CXCR4 or SNAP-CXCR4. (a) Pre-incubation with VUN400-HiBiT; ligands were added following a 30-minute incubation with VUN400-HiBiT. Cells were subsequently incubated for two hours at 37°C and luminescence measured. (b) Simultaneous addition; ligands were added at the same time as VUN400-HiBiT. Cells were subsequently incubated for two hours at 37°C and luminescence measured. (c) Pre-incubation with ligands; ligands were added and the cells were subsequently incubated for two hours at 37°C prior to addition of 20 nM VUN400-HiBiT. Cells were then incubated for a further 30 minutes before luminescence was measured. Data are mean ± SEM from n repeats. *p<0.05, unpaired t-test comparing pIC_50_ at SNAP-CXCR4 to that at LgBiT-CXCR4. ^†^p<0.05, unpaired t-test comparing pIC_50_ to that obtained when pre-treating tagged CXCR4 with VUN400-HiBiT. ^#^p<0.05, unpaired t-test comparing maximal inhibition of VUN400-HiBiT binding to 10 µM AMD3100.

| Ligand | LgBiT-CXCR4 |  | n | SNAP-CXCR4 |  | n |
| --- | --- | --- | --- | --- | --- | --- |
|  | pIC<sub>50</sub><br>(mean ± SEM) | Max inhibition<br>(% 10 µM<br>AMD3100) |  | pIC<sub>50</sub><br>(mean ± SEM) | Max inhibition<br>(% 10 µM<br>AMD3100) |  |
| *(a) Pre-incubation with VUN400-HiBiT* |  |  |  |  |  |  |
| CXCL12 | 8.82 ± 0.12 | 98.2 ± 0.2 | 5 | 8.14 ± 0.28* | 93.3 ± 8.7 | 3 |
| AMD3100 | 7.13 ± 0.18 | 100 | 5 | 6.94 ± 0.16 | 100 | 3 |
| IT1t | 6.93 ± 0.09 | 104.4 ± 4.5 | 5 | 7.36 ± 0.12 | 107.1 ± 4.8 | 3 |
| *(b) Simultaneous addition* |  |  |  |  |  |  |
| CXCL12 | 8.74 ± 0.13 | 98.8 ± 0.3 | 6 | 8.05 ± 0.21* | 92.8 ± 6.7 | 4 |
| AMD3100 | 6.69 ± 0.19 | 100 | 5 | 7.03 ± 0.24 | 100 | 8 |
| IT1t | 7.29 ± 0.04 | 96.5 ± 0.4 | 3 | 7.53 ± 0.08 | 106.8 ± 11.5 | 3 |
| *(c) Pre-incubation with ligands* |  |  |  |  |  |  |
| CXCL12 | 9.03 ± 0.02 | 111.5 ± 3.1<sup>#</sup> | 6 | 8.05 ± 0.41 | 67.7 ± 11.6<sup>#</sup> | 6 |
| AMD3100 | 7.93 ± 0.16<sup>†</sup> | 100 | 6 | 7.78 ± 0.24<sup>†</sup> | 100 | 7 |
| IT1t | 7.68 ± 0.13<sup>†</sup> | 79.9 ± 3.1<sup>#</sup> | 5 | 7.52 ± 0.13 | 102.3 ± 3.6 | 7 |

**Figure 5.**
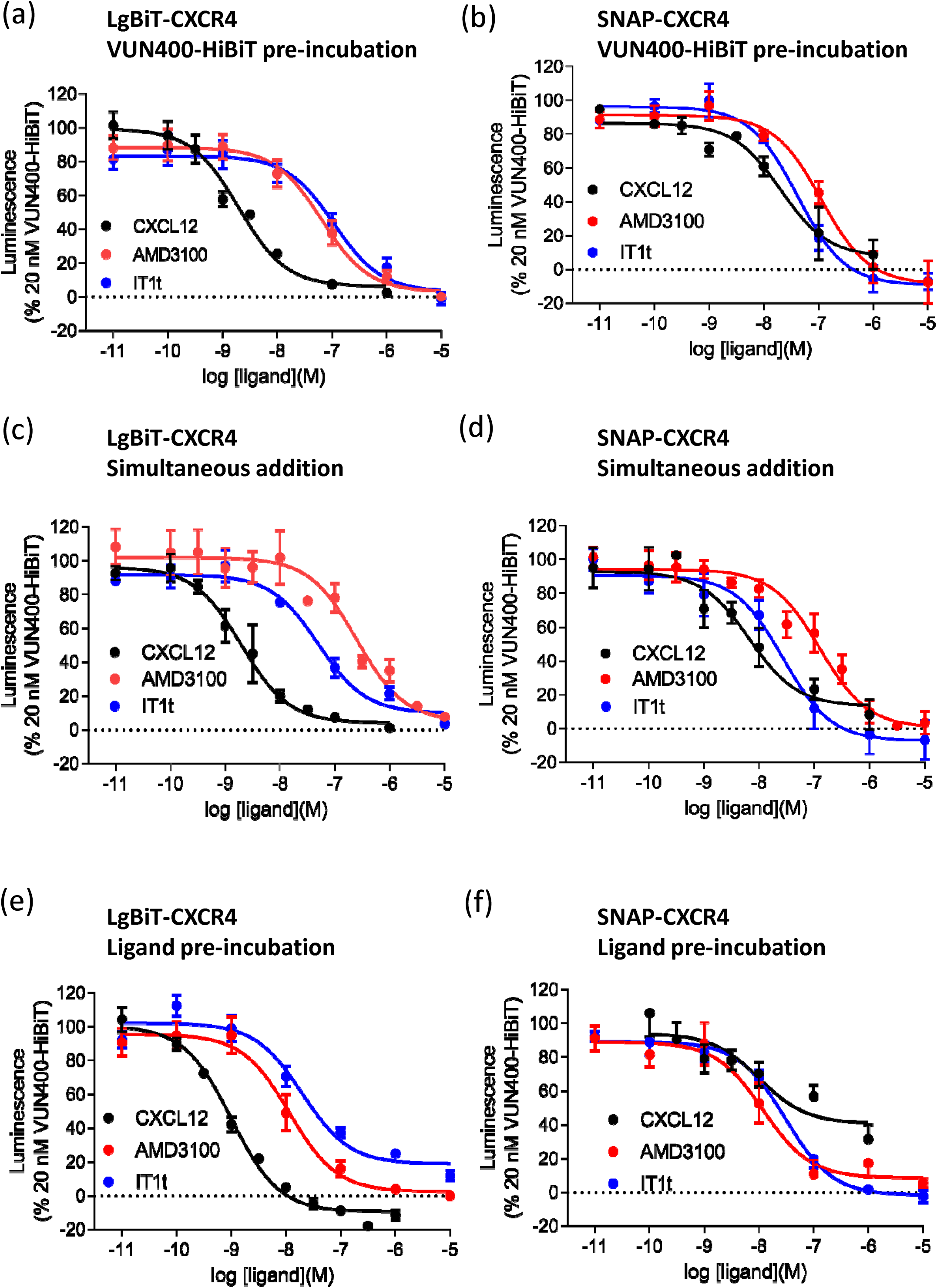
Displacement of VUN400-HiBiT by competing ligands. HEK293 cells stably expressing LgBiT-CXCR4 (a,c,e) or SNAP-CXCR4 (b,d,f) were treated with 20 nM VUN400-HiBiT and CXCL12, AMD3100, IT1t. (b,d,f) 20 nM exogenous LgBiT added to complement with bound VUN400-HiBiT. (a,b) Cells treated with 20 nM VUN400-HiBiT for 30 minutes prior to the addition of CXCL12, AMD3100 or IT1t. (c,d) Cells treated with VUN400-HiBiT and CXCL12, AMD3100 or IT1t simultaneously. (e,f) Cells treated with CXCL12, AMD3100 or IT1t for two hours prior to the addition of VUN400-HiBiT. Data are combined mean ± SEM from at least three separate experiments, where each experiment was performed in triplicate. See Table 1 for full details on number of experimental repeats.

To determine whether the reduction in bioluminescence upon AMD3100 or IT1t treatment was due to altered bioluminescence complementation (e.g. a change in orientation rather than displacement of VUN400-HiBiT from the receptor), we investigated the binding of the VUN400-HiBiT to CXCR4 with a N-terminal SNAP-tag. As a 19 kDa protein, the SNAP-tag is a similar in size to the full length NanoLuc, thus acting as a steric N-terminal control for HiBiT-LgBiT complementation. Additionally, the SNAP-tag was not luminescent. Therefore, exogenous LgBiT (20 nM) was added at the end of the experiment, in order to complement with VUN400-HiBiT bound to SNAP-CXCR4. VUN400-HiBiT binding to the SNAP-CXCR4 receptor was observed (Supplementary Figure 1). However, binding was linear over the concentration of 1-500nM, indicative of lower affinity binding. The expression level (as determined by the overall luminescence achieved) also appeared to be much lower than that obtained with LgBiT-CXCR4 (Supplementary Figure 1 and Figure 2). Nevertheless, AMD3100 was still able to reduce the bioluminescence signals. Because complementation was achieved with LgBiT in solution, it was unlikely that the reduced signals by AMD3100 were caused by changes in HiBiT-LgBiT proximity. Therefore, these data suggest that AMD3100 causes a reduction in signals by reducing the binding of VUN400-HiBiT to CXCR4.

The binding of VUN400-HiBiT to SNAP-CXCR4 was inhibited by CXCL12, AMD3100 and IT1t in a concentration-dependent manner (Figure 5b; Table 1a). The resulting IC_50_ values of AMD3100 and IT1t determined at the SNAP-CXCR4 were not significantly different from those measured at the LgBiT-CXCR4 (Table 1a, p>0.05, unpaired t-test), suggesting that NanoBiT complementation between the VUN400-HiBiT and LgBiT-CXCR4 had no effect on the ability of these antagonist ligands to prevent binding of the nanobody to CXCR4. Interestingly, CXCL12 had a lower pIC_50_ value in the SNAP-CXCR4 cell line than in the LgBiT-CXCR4 cell line (p<0.05, unpaired t-test).

Next, the effects of simultaneous addition of VUN400-HiBiT and ligands were investigated. Whole cells were treated with 20 nM VUN400-HiBiT, which was added to the plate simultaneously with increasing concentrations of the endogenous agonist CXCL12, or the antagonists AMD3100 and IT1t. Figure 5c shows a clear decrease in luminescence at the LgBiT-CXCR4 in the presence of high concentrations of competing ligands (Table 1b). Similar results were obtained at the SNAP-CXCR4 receptor (Figure 5d, Table 1b). Again, CXCL12 had a lower pIC_50_ value in the SNAP-CXCR4 cell line than in the LgBiT-CXCR4 cell line (p<0.05, unpaired t-test). Taken together, these data show that the binding of CXCL12, AMD3100 and IT1t to N-terminally tagged CXCR4 receptors can inhibit VUN400-HiBiT binding to ECL2 of the receptor, probably as a result of a conformational change. This conformational change may also change the orientation of the HiBiT on VUN400-HiBiT bound to ECL2 with respect to LgBiT on the N-terminus LgBiT-CXCR4 (effectively preventing complementation), allowing more sensitive detection of agonist-induced conformational changes with the NanoLuciferase complementation approach than with traditional ligand binding approaches (Figure 5).

### Conformational selectivity of VUN400-HiBiT

The VUN400 nanobody was originally selected using phage display, utilising CXCR4-expressing lipoparticles containing high concentrations of CXCR4 in native conditions, and thus in a myriad of specific conformations. Therefore, there was the potential that VUN400 recognised a specific CXCR4 conformation which could be modulated by ligand treatment. With this in mind, we repeated the VUN400-HiBiT displacement experiments at the LgBiT-CXCR4, incubating with CXCL12, AMD3100 or IT1t prior to the addition of VUN400-HiBiT. The binding of VUN400-HiBiT to LgBiT-CXCR4 was inhibited by all ligands with a higher potency than when ligands and nanobody were added simultaneously (Figure 5e, Table 1c). This effect was particularly marked with AMD3100, where the potency was an order of magnitude higher when cells were pre-treated with AMD3100 for 2h prior to addition of VUN400-HiBiT (Figure 5e; Table 1c). These data suggest that AMD3100 (and to a lesser extent IT1t) can induce a conformation that has higher affinity for the small molecule than when the nanobody is added at the same time. This is suggestive of a negative allosteric interaction between the binding sites for AMD3100 and VUN400-HiBiT, but could also be due to the ECL2 bound nanobody sterically interfering with access of the small molecule inhibitor to its transmembrane binding site. A similar observation was made for AMD3100 with the SNAP-CXCR4 receptor (Figure 5e; Table 1c).

Interestingly, in the case of CXCL12 binding to SNAP-CXCR4, the inhibition curve appeared to be much less potent that in the equivalent LgBiT-CXCR4 experiments and the plateau above zero indicative of the presence of multiple components of VUN400-HiBiT binding; one of which was insensitive to displacement by low concentrations of CXCL12 (Figure 5f). This was not a result of the N-terminal SNAP-tag interfering with the binding of CXCL12 to SNAP-CXCR4 (Supplementary Figure 2). TR-FRET ligand binding confirmed an affinity of CXCL12-AF647 (pK_D_ 7.99 ± 0.10; n=5, Supplementary Figure 2) that was in agreement with that measured at the LgBiT-CXCR4 with NanoBRET.

To determine the temporal characteristics of any conformational changes induced by AMD3100 and CXCL12, LgBiT-CXCR4 receptors were treated with VUN400-HiBiT, allowed to reach equilibrium, and then competing with either CXCL12 or AMD3100. The addition of these ligands resulted in a decrease in luminescence over time, signifying reduced complementation of VUN400-HiBiT with LgBiT-CXCR4 (Figure 6a). Fitting a single exponential curve to the data revealed apparent *k*_*off*_ rates for VUN400-HiBiT when either AMD3100 (*k*_*off*_ 0.039 ± 0.002 min^−1^; n=4) or CXCL12 (*k*_*off*_ 0.046 ± 0.001 min^−1^; n=5) were simultaneously present (p<0.05 comparing *k*_*off*_ values, unpaired t-test). In comparison, the fluorescent ligand CXCL12-AF647 showed much faster kinetic off rates in the presence of both ligands with no statistical difference between the *k*_*off*_ values (CXCL12 *k*_*off*_ 0.32 ± 0.03 min^−1^; AMD3100 *k*_*off*_ 0.42 ± 0.04 min^−1^; n=4, Figure 6b).

**Figure 6.**
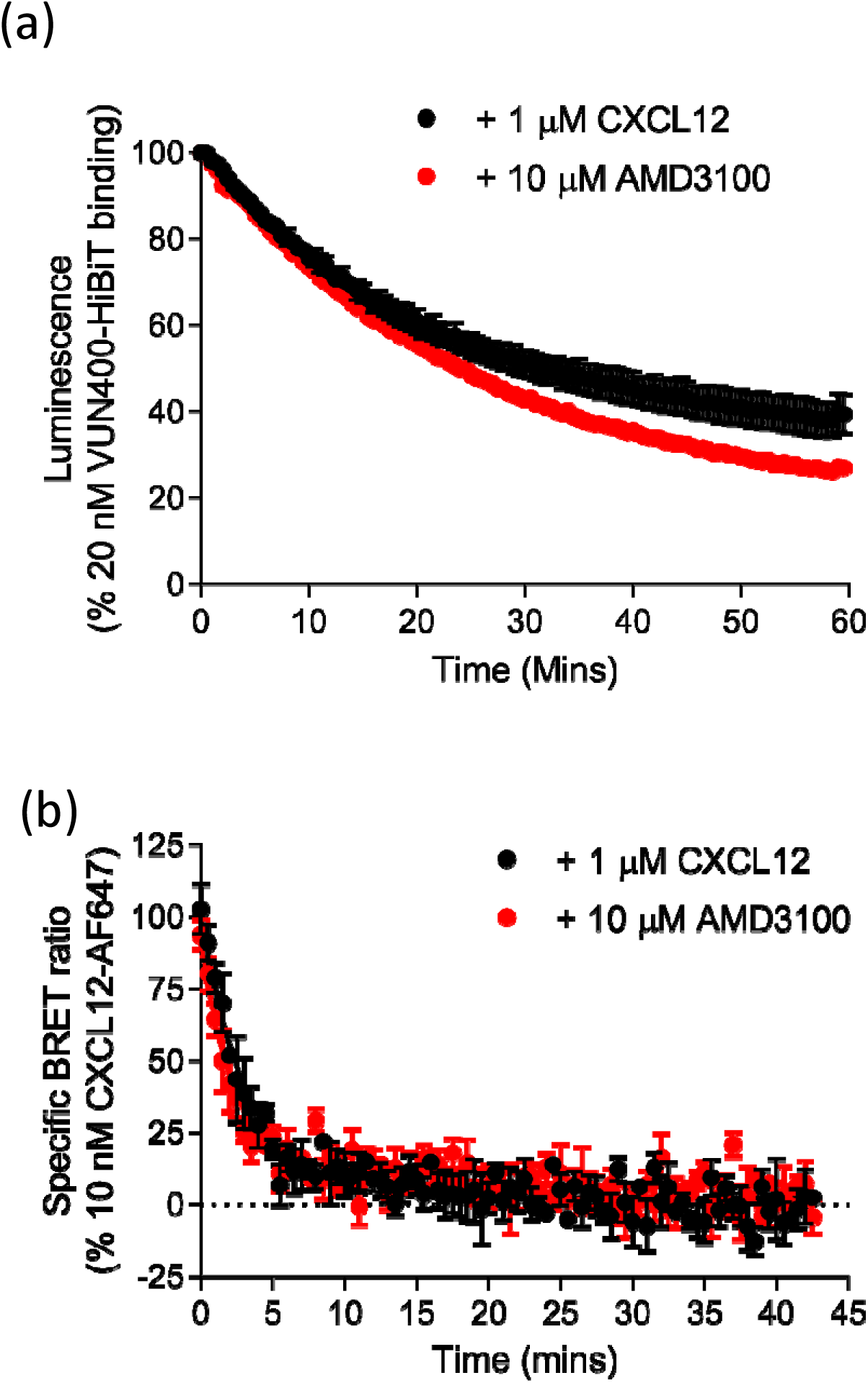
Dissociation kinetics of probe-receptor complex. (a) Decrease in VUN400-HiBiT-LgBiT-CXCR4 complemented luminescence over time following addition of 1 µM CXCL12 or 10 µM AMD3100. (b) Decrease in specific BRET ratio from 10 nM CXCL12-AF647 following the addition of 1 µM CXCL12 or 10 µM AMD3100. Data are combined mean ± SEM from (a -CXCL12) five or (a - AMD3100; b) four separate experiments, where each experiment was performed in triplicate.

## Discussion

Nanobodies have been used previously to stabilise GPCRs and investigate membrane protein conformations to elucidate new aspects of GPCR function (Zimmermann et al., 2018; Heukers et al., 2018; De Groof et al., 2019a; Heukers et al., 2019). For example, nanobodies recognizing active GPCRs have been used to monitor G protein-mediated signalling after receptor internalization (Irranejad et al., 2013, Stoeber et al., 2018). The ability to directly measure nanobody-target engagement is crucial to understand the nature of this interaction. Split NanoLuciferase technology (NanoBiT) offered the possibility to use complemented luminescence to measure small changes in the interactions between proteins (nanobody-receptor in this case). This high-throughput, plate-reader-based approach allowed for the continuous detection of luminescence to measure nanobody affinity and kinetic rate constants of nanobody binding to receptors in a living cell environment.

The expression and ligand binding capabilities of the LgBiT-tagged CXCR4 were first confirmed in HEK293 cells. LgBiT-tagged CXCR4 was able to successfully complement with exogenously applied HiBiT-HaloTag to form the full-length NanoLuc luciferase and cell surface luminescence was detected using plate-reader and bioluminescence imaging modalities. Since the size of the HiBiT-HaloTag fusion protein (34 kDa) prevented it from crossing the plasma membrane, the complemented luminescence was effectively confined to receptors expressed on the cell surface. The resulting affinity of purified HiBiT for LgBiT-CXCR4 (pK_D_ 6.43) was significantly lower than those determined using purified HiBiT and LgBiT fragments (pK_D_ 9.15; Dixon et al., 2016). We have similarly shown that HiBiT-CXCR4 also has a much lower affinity for purified LgBiT (pK_D_ 6.64; White et al., 2020). It is therefore likely that this is a consequence of steric hindrance caused by the extracellular components of the receptor to which the NanoBiT tag is attached. The similarity between the pK_D_ values obtained here for purified HiBiT-HaloTag interaction with LgBiT-CXCR4 (6.4) and those obtained with HiBiT-CXCR4 and purified LgBiT (White et al., 2020) suggest that this is not a consequence of the HaloTag component of the purified HiBiT used here. Consequently, the kinetic values obtained in this study are considered to be a relatively accurate representation of the binding kinetics of the nanobody to CXCR4.

NanoBRET ligand binding also confirmed that the receptor could tolerate the addition LgBiT via N-terminal fusion without the loss of binding affinity of the fluorescent ligand CXCL12-AF647. This tolerance is in agreement with previous studies on NanoBRET-based ligand binding with CXCR4 receptor tagged on the N-terminus with full-length NanoLuc (White et al., 2020). Therefore, even though the N-terminus of CXCR4 is involved in the binding of CXCL12 (Crump et al., 2018; Gupta et al., 2001), our NanoBRET binding data showed that N-terminally fused LgBiT did not interfere with ligand binding. Similarly, the SNAP tag on the N-terminus of the receptor did not interfere with the binding of CXCL12 to SNAP-CXCR4. In our studies, the affinity of CXCL12-AF647 measured using TR-FRET was the same as that determined at the LgBiT-CXCR4.

Single domain antibodies represent powerful tools for interrogating GPCR pharmacology. In most of the studies that employed nanobodies to elucidate GPCR structural and functional information, intracellularly binding, non-modified nanobodies have been used (Rasmussen et al., 2011; Huang et al., 2015; Che et al., 2018). Here, the CXCR4-targeted nanobody VUN400 was modified on its C-terminal tail to incorporate the HiBiT polypeptide tag.

Nanobody binding to LgBiT-CXCR4 was subsequently detected using complemented luminescence. The resulting affinity of VUN400-HiBiT at LgBiT-CXCR4 (pK_D_ 7.25) was in good agreement with previous data determining the affinity of VUN400 at the unmodified CXCR4 receptor with radioligand binding (pIC_50_ 7.3; Bobkov et al., 2018). These data are also in agreement with the general observation that C-terminal fusion to nanobodies does not affect their binding characteristics (Fang et al., 2016). Kinetic measurements of nanobody-receptor interactions have traditionally been performed using purified receptors with surface plasmon resonance assays, often in an environment which is not indicative of the natural environment of the cell. Our approach allowed for the measurement of kinetic rate constants of nanobody-receptor binding (*k*_*on*_ 1.98×10^5^ M^−1^ min^−1^, *k*_*off*_ 0.032 min^−1^) in living cells. Taken together, these data strongly suggested that the measured affinity and kinetic rate constants were a result of nanobody-receptor binding, rather than HiBiT-LgBiT complementation.

VUN400-HiBiT could also detect SNAP-tagged CXCR4 receptors without the presence of the LgBiT tag fused to the receptor, albeit with a reduced apparent affinity. The apparent lower affinity of VUN400-HiBiT for SNAP-CXCR4 is likely to be due to steric hindrance because of the need for the added purified LgBiT fragment to also access the HiBiT peptide sequence to obtain the final bioluminescence signal. More interestingly, VUN400-HiBiT was able to label wild-type CXCR4 receptors expressed under endogenous promotion in Jurkat cells following addition of exogenous purified LgBiT. This makes this technology applicable for assessing endogenous CXCR4 expression and conformation on a wide variety of cell lines or tissues.

Interestingly, the addition of saturating concentrations of chemokine was able to prevent complementation between VUN400-HiBiT and LgBiT-CXCR4 in transfected HEK cells. This was also seen in Jurkat cells endogenously expressing wild-type CXCR4. This suggested that NanoBiT complementation was reversible, which is consistent with the observed lower affinity of purified HiBiT for LgBiT-CXCR4. The mechanisms by which CXCL12 and small molecule antagonists, such as AMD3100, displace VUN400-HiBiT binding ECL2 of CXCR4 is likely to be allosteric and a consequence of conformational changes induced in the receptor structure, since the N-terminus of CXCR4 is involved in the binding of CXCL12 (Crump et al., 2018; Gupta et al., 2001) and small molecules such as AMD3100 bind to the transmembrane regions of CXCR4 (Rosenkilde et al 2004). Furthermore, there is no overlap between the binding site of VUN400 in the ECL2 of CXCR4 (Van Hout et al., 2018), and the binding sites of the small molecule inhibitors AMD3100 (Rosenkilde et al., 2004) or IT1t (Wu et al., 2010), which are within the upper transmembrane region of the receptor.

In all experimental conditions, increasing concentrations of ligands resulted in a loss of VUN400-HiBiT binding to LgBiT-CXCR4 (Figure 5). Adding VUN400-HiBiT before competing ligands gave similar IC_50_ values to those determined when VUN400-HiBiT was added simultaneously to the ligands. However, when competing ligands were allowed to bind to LgBiT-CXCR4 before being challenged with VUN400-HiBiT, there was an apparent increase in the affinities (reduction in IC_50_ value) of these ligands (Table 1c). This was particularly marked for AMD3100, where the potency was an order of magnitude higher when cells were pre-treated with AMD3100 for 2h prior to addition of VUN400-HiBiT. This change in IC_50_ value is likely to be due to the nanobody bound to ECL2 sterically interfering with subsequent access of the small molecule inhibitor to its transmembrane binding site when both are present together.

Taken together, these data show that the binding of AMD3100 and IT1t to N-terminally tagged CXCR4 receptors can inhibit VUN400-HiBiT binding to ECL2 of the receptor, probably as a result of a conformational change. This is consistent with a reciprocal negative allosteric interaction between the binding sites for AMD3100 and VUN400-HiBiT, and the known ability of small molecule inhibitors such as AMD3100 to induce conformational rearrangement of the extracellular domains of CXCR4 (White et al., 2020) and modulate monoclonal antibody binding (Carnec et al., 2005; Rosenkilde et al., 2004). The conformational changes involving ECL2 induced by both CXCL12 and AMD3100 in LgBiT-CXCR4 could be followed in real-time in living cells following pre-equilibration with VUN400-HiBiT bound (Figure 6) and these changes were much slower than the ability of both ligands to induce dissociation of a fluorescent analogue of CXCL12 from the receptor. It is well known that CXCR4 forms dimers or oligomeric complexes (Percherancier et al., 2005; Wang et al., 2006), and that this organization into higher order structures can affect CXCR4 function (Lagane et al., 2008; Ge et al., 2017). It remains to be established whether the conformational changes induced by CXCL12 or AMD3100 that are influencing the binding of VUN400-HiBiT to CXCR4 involve monomeric species of CXCR4 or higher order oligomeric receptor species.

A striking feature of the effect of CXCL12 on VUN400-HiBiT binding is the difference in CXCL12-sensitivity between VUN400-HiBiT binding to LgBiT-CXCR4 and SNAP-CXCR4. In contrast, the small molecule inhibitors were equally effective in inhibiting VUN400-HiBiT binding to LgBiT-CXCR4 and SNAP-CXCR4. It is possible that this is not a difference in binding but rather in NanoLuciferase complementation. The conformational change induced by the agonist CXCL12 may affect the relative orientation of the HiBiT on VUN400-HiBiT (attached to ECL2) towards the LgBiT on the N-terminus of CXCR4 (effectively preventing complementation). This could allow a more sensitive detection of agonist-induced conformational changes by NanoLuciferase complementation (Figure 5).

In conclusion, the data presented here demonstrate the ability to detect ligand binding in living cells to CXCR4 using NanoBiT complementation technology in combination with an extracellularly binding, ECL2-targeted, nanobody. Monitoring the complemented luminescence enabled the kinetics of the nanobody-receptor interaction to be followed in real-time, as well as probing for ligand-induced conformational changes in the extracellular regions of CXCR4. Furthermore, the unique selectivity of VUN400-HiBiT, and the exquisite sensitivity of NanoLuciferase bioluminescence, enabled its use to monitor ligand-binding to and ligand-induced conformational changes of the endogenous wild-type CXCR4 on the cell surface.

## Significance

The chemokine receptor CXCR4 is a G protein-coupled receptor that plays an important role in the immune response and in the progression of many diseases, including cancer and HIV infection. Camelid single-domain antibody fragments (nanobodies) offer the specificity of an antibody in a single 15kDa immunoglobulin domain. Their small size allows for easy genetic manipulation of the nanobody sequence to incorporate protein tags, facilitating their use as biochemical probes. Here, we have used the nanobody VUN400, which recognises the second extracellular loop (ECL2) of the human CXCR4, and NanoBiT luciferase complementation to measure the direct binding of VUN400-HiBiT to the second extracellular loop of CXCR4 receptor, tagged with LgBiT, in living cells and in real time. We also show that this approach can be used to monitor conformational changes in the extracellular domains of the receptor induced by agonists and antagonists in living cells. For example, the clinically-used small molecule CXCR4 antagonist AMD3100 was able to displace VUN400-HiBiT binding to ECL2 of CXCR4, despite binding to the transmembrane regions of the receptor. VUN400-HiBiT could also detect cell surface expression of recombinant CXCR4 not containing the LgBiT tag as well as wild-type receptors endogenously expressed in Jurkat T-cells. This ability to probe for endogenous receptor expression and ligand-induced conformation changes offers a powerful tool to investigate G protein-coupled receptor function in healthy and diseased cells.

## Footnote

Part of this work was published in abstract form as part of the conference proceedings of Experimental Biology 2019 (Soave et al., 2019a).

## Acknowledgements

We thank Timo De Groof (Vrije Universiteit Amsterdam, The Netherlands) for his assistance with VUN400-HiBiT nanobody purification. This work was supported by MRC (grant number MR/N020081/1; M.S., B.K., J.W., S.J.B., and S.J.H.), the European Union (H2020-MSCA program grant agreements 641833-ONCORNET [M.J.S., S.J.B., and S.J.H.] and 860229-ONCORNET 2.0; [R.H., M.J.S., S.J.B., and S.J.H.]), and the Dutch Research Council (ENPPS.TA.019.003 MAGNETIC; R.H. and M.J.S).

## Author Contributions

M.S. performed the experiments, analyzed data, developed the NanoBiT-based nanobody binding protocol, and wrote the manuscript. R.H. helped the production of the VUN400 nanobody, coordinated the project, and wrote the manuscript. B.K. and J.W. supervised the project and wrote the manuscript. M.J.S. coordinated the project and wrote the manuscript. S.J.B. conceived, coordinated and supervised the project, and wrote the manuscript. S.J.H. conceived, coordinated and supervised the project, and wrote the manuscript.

## Competing Interests

R.H. is CSO of QVQ Holding B.V.

## STAR Methods

### Experimental Model and Subject Details

Jurkat (male) and HEK293 GloSensor (female) cells were transfected and cultured as described in Method Details.

### Method Details

#### Materials

BL21 (DH3) competent cells were obtained from Agilent Technologies (Santa Clara, CA, USA). Isopropyl-β-D-thiogalactopyranoside and AMD3100 were purchased from Sigma Aldrich. All restriction enzymes, purified LgBiT, purified HiBiT, FuGENE® transfection reagent, and furimazine were purchased from Promega (Wisconsin, USA). Phosphate buffered saline (PBS), Dulbecco’s modified Eagle’s medium (DMEM), foetal calf serum (FCS), and bovine serum albumin (BSA) were obtained from Sigma Aldrich (UK). CXCL12 was purchased from PeproTech. IT1t was purchased from Tocris Bioscience. Fluorescent CXCL12-AF647 was obtained from Almac (Craigavon, UK).

#### DNA constructs

To create the LgBiT-tagged human CXCR4 receptor construct (LgBiT-CXCR4), the LgBiT sequence was amplified from the pBiT1.1-N [TK/LgBiT] vector (Promega) and assembled in-frame with the membrane sequence of the 5-HT_3A_ membrane localisation signal sequence (sig) at the 5’ end of LgBiT using the Gibson assembly PCR technique. The 3’ end of LgBiT was modified to incorporate a short linker (GSSG) and the recognition site for the tobacco etch virus (TEV) enzyme (residue sequence: EDLYFQS). The resulting nucleotide fragment (sig.LgBiT) contained the sequences for the restriction enzyme KpnI upstream of the sig sequence and the restriction enzyme BamHI downstream of the linker. This was then ligated to the pcDNA3.1 plasmid containing the human CXCR4 receptor (described in Adlere et al., 2019), creating the fusion of sig.LgBiT, a Gly-Ser linker and CXCR4 with the methionine start signal removed.

#### Cell Culture

Jurkat cells (clone E6-1) were maintained in T175 flasks containing RPMI 1640 medium (Lonza) supplemented with 10% fetal calf serum (FCS; Sigma Aldrich) and 2 mM L-glutamine (Sigma Aldrich) at 37°C/5% CO_2_. Fresh medium was added to the cells every 2-3 days. Cells were passaged at 70% confluency by withdrawing 2.5 ml of the cells into a fresh T175 flask with medium. HEK293G cells (Glosensor cAMP HEK293, Promega) were grown in T75 flasks containing 25 ml Dulbecco’s Modified Eagle’s Medium (DMEM; Sigma Aldrich, UK) supplemented with 10% FCS at 37°C/5% CO_2_. Cells were passaged at 70-80% confluency using phosphate buffered saline (PBS; Lonza) and trypsin (0.25% w/v in versene; Lonza). A mixed population HEK293G cell line was created by transfecting cells with the LgBiT-CXCR4 construct using FuGENE® (Promega) according to the manufacturer’s instructions, followed by selective pressure (1 mg/ml G418) for two to three weeks. HEK293G cells stably expressing the SNAP-CXCR4 construct were kindly gifted from Dr. J. Goulding (University of Nottingham).

#### Generation and purification of VUN400-HiBiT nanobody

The pET28a bacterial expression vector containing the monovalent VUN400 nanobody (Bobkov et al., 2018; Van Hout et al., 2018) was modified at the C-terminus by fusing the in frame sequence of the HiBiT peptide tag (VSGWRLFKKIS) in between the C-terminal cMyc and His-tags using PCR cloning, generating the VUN400-HiBiT construct. The pET28a VUN400-HiBiT plasmid was transformed into BL21 (DH3) *Escherichia coli* cells (Agilent Technologies) for expression and purification. VUN400-HiBiT was then purified as previously described (Wit et al., 2010). Briefly, periplasmic expression of VUN400-HiBiT was induced by 1 mM isopropyl-β-D-thiogalactopyranoside to the bacterial culture media. Periplasmic extracts were obtained from the freezing and thawing of the cell pellet, and were resuspended in PBS. VUN400-HiBiT was purified from the periplasmic extract by ion-metal affinity chromatography using Ni-NTA agarose resin (ThermoFischer) according to the manufacturer’s instructions. Finally, a buffer exchange to PBS was performed using SnakeSkin (ThermoFischer) dialysis.

#### NanoBRET Binding Assay

Saturation NanoBRET binding assays were performed on HEK 293 cells stably expressing the LgBiT-CXCR4 receptor. Cells were plated at 30,000 cells/well in white 96-well plates (Greiner) were coated with 50 µl poly-D-lysine (mol wt: 70-150 kDa; 10 µg/ml in PBS) in 100 µl complete media (DMEM with 10% FCS). After 24 hours, the media was removed from each well and replaced with 100 µl HEPES-buffered saline solution (HBSS; 145 mM NaCl, 5 mM KCl, 1.3 mM CaCl_2_, 1 mM MgSO_4_, 10 mM HEPES, 2 mM sodium pyruvate, 1.5 mM NaHCO_3_, 10 mM D-glucose, pH 7.45) containing 10 nM purified HiBiT-HaloTag (Promega) and incubated at 37°C for 30 minutes. This was sufficient time for HiBiT-LgBiT complementation to occur, forming the fully complemented NanoLuc enzyme. The HBSS was then aspirated and the cells washed three times with 100 µl HBSS. Cells were then treated with 100 µl HBSS containing the relevant concentration of fluorescent ligand in the absence or presence of 10 µM AMD3100 to define non-specific binding. Cells were incubated for 2 hours at 37°C in the dark. The NanoLuc substrate furimazine (Promega) was added to each well (1:400 final concentration) and allowed to equilibrate for 5 minutes at 37°C prior to reading. Luminescence signals were measured at two wavelengths using a PHERAstar FS plate reader (BMG Labtech, UK) at room temperature. Filtered light was simultaneously measured using 460 nm (80-nm bandpass) and >610 nm longpass filters. The resulting BRET ratio was calculated by dividing the >610 nm emission by the 460 nm emission.

#### Bioluminescence Imaging

Bioluminescence imaging was performed with an Olympus LV200 Widefield inverted microscope, equipped with a 60x/1.42 NA oil immersion objective lens. HEK293G cells stably expressing the LgBiT-CXCR4 construct were seeded on a poly-D-lysine coated (10 µg/ml) 35mm MatTek dish with a No 1.5 glass coverslip at a density of 120,000 cells per dish and incubated in complete media at 37°C in a humidified 95% air/5% CO_2_ environment overnight. Prior to imaging, media was removed and replaced with 2 ml HBSS and the dish was placed on the heated stage of the LV200 for 15 minutes at 37°C. For LgBiT-CXCR4 imaging, 200 µl HBSS containing furimazine (1:400 final concentration) and either 20 nM purified HiBiT-HaloTag control protein (Promega) or 20 nM VUN400-HiBiT was added to the dish, and the subsequent luminescence was captured in the open channel with a 45 second (+ 20 nM HiBiT-HaloTag) or 10 second (+ 20 nM VUN400-HiBiT) exposure using a 0.688 MHz EMCCD with a gain of 200.

#### NanoBiT nanobody saturation binding assay

Saturation binding with VUN400-HiBiT was performed on HEK 293 cells stably expressing the LgBiT-CXCR4 receptor. Cells were plated at 30,000 cells/well in white 96-well plates were coated with poly-D-lysine (10 µg/ml in PBS) in 100 µl complete media (DMEM with 10% FCS). After 24 hours, the media was removed and replaced with 100 µl HBSS containing VUN400-HiBiT within the range of 4-500 nM. Where used, CXCL12, AMD3100 or IT1t were also added at this point. The plate was incubated for two hours at 37°C. Furimazine (1:400 final concentration) was added to each well and the plate was incubated for a further 15 minutes at 37°C. Luminescence was measured using the PHERAstar FS plate reader.

#### NanoBiT nanobody displacement binding assay

Full nanobody displacement assays were performed in HEK293 cells stably expressing either the LgBiT-CXCR4 or SNAP-CXCR4 constructs. Cells (30,000 per well) were plated in white 96-well poly-D-lysine coated plates as described above. After 24 hours, media was removed and replaced with 100 µl HBSS. Increasing concentrations of displacing ligands were added to the cells. Additionally, cells were treated with 20 nM VUN400-HiBiT. Depending on the experimental condition, this was as a 30 minute pre-treatment prior to ligand treatment, simultaneously with competing ligands, or 2 hours following ligand treatment. The cells were incubated at 37°C for 2 and a half hours in total. SNAP-CXCR4 cells were washed 3x with 100 µl warm HBSS to remove unbound VUN400-HiBiT and then treated with 20 nM exogenous LgBiT to complement with bound VUN400-HiBiT. Furimazine (1:400 final concentration) was added to each well and the plate was incubated for a further 15 minutes at 37°C. Luminescence was measured using the PHERAstar FS plate reader.

Nanobody displacement assays in T-cells were performed by seeding 100,000 Jurkat cells per well into V-bottom 96-well plates in 100µl warm HBSS. Cells were treated with 100 nM VUN400-HiBiT and competing ligands simultaneously and incubated for 2 hours at 37°C. The cells were pelleted by centrifugation at 300*g* for 5 minutes and the supernatant removed and replaced with 200 µl fresh HBSS. Cells were resuspended via shaking at 1000 r.p.m on an orbital plate shaker for 2 minutes. This wash step was repeated two more times to remove unbound VUN400-HiBiT. The cells were treated with 20 nM exogenous LgBiT and furimazine (1:400 final concentration) and transferred to white 96-well plates and luminescence read using a PHERAstar FS plate reader.

#### NanoBiT kinetic measurements

Association kinetic measurements of HiBiT or nanobody binding were performed in HEK293 cells stably expressing LgBiT-CXCR4. Cells (30,000 per well) were plated in white 96-well poly-D-lysine coated plates (10 µg/ml in PBS) as described above. After 24 hours, media was removed and replaced with 50 µl HBSS containing furimazine (1:400 final concentration) and incubated at 37°C for 10 minutes and basal luminescence was measured. Increasing concentrations of HiBiT or VUN400-HiBiT were added to the cells in 50 µl volumes per well. The resulting luminescence was measured over 60 minutes at 37°C. Luminescence was measured using the PHERAstar FS plate reader.

Nanobody and CXCL12-AF647 dissociation kinetic measurements were performed in membranes isolated from HEK293 LgBiT-CXCR4 cells. Briefly, 10 µg LgBiT-CXCR4 membranes were plated in white 96-well plates containing furimazine (1:400 final concentration) and incubated at 37°C for 10 minutes. For experiments with CXCL12-AF647, LgBiT-CXCR4 membranes were incubated with 20 nM purified HiBiT during this time to ensure complete complementation. Increasing concentrations of VUN400-HiBiT or CXCL12-AF647 were added to the wells and luminescence (VUN400-HiBiT) or the BRET ratio (CXCL12-AF647) were measured. A saturating concentration of competing ligands (CXCL12 or AMD3100) were added following a 30 minute (VUN400-HiBiT) or 15 minute (CXCL12-AF647) incubation and the luminescence or BRET ratio was measured for another hour. The BRET ratio was calculated as described above. Both the BRET ratio and luminescence were measured using the PHERAstar FS plate reader.

#### Data analysis

Data were analysed using Prism 7 software (GraphPad, San Diego, USA).

Saturation NanoBRET curves were fitted simultaneously for total (CXCL12-AF647) and non-specific binding (in the presence of 10µM AMD3100) using the following equation:

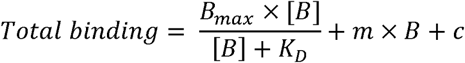

where *B*_*max*_ is the maximal specific binding, [B] is the concentration of the fluorescent ligand (nM), *K*_*D*_ is the equilibrium dissociation constant (nM), *m* is the slope of the non-specific binding component, and c is the y-axis intercept.

Saturation binding curves of VUN400-HiBiT were fitted using the following equation:

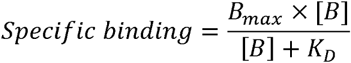

where *B*_*max*_ is the maximal specific binding, [B] is the concentration of VUN400-HiBiT, and *K*_*D*_ is the equilibrium dissociation constant (nM).

The affinities of ligands at the LgBiT-CXCR4 and SNAP-CXCR4 receptors were calculated from VUN400-HiBiT binding data with a one-site sigmoidal response curve given by the following equation:

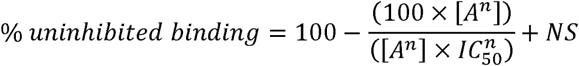

where [A] is the concentration of unlabelled ligand, *NS* is non-specific binding, *n* is the Hill coefficient, and *IC*_*50*_ is the concentration of ligand required to inhibit 50% of VUN400-HiBiT.

Nanobody binding association kinetic data were fitted to the following mono-exponential association function:

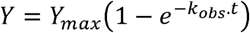

where Y_max_ is the specific binding at infinite time, *t* is the time of incubation, and *k*_*obs*_ is the rate constant for the observed rate of association.

*k*_*on*_ and *k*_*off*_ values were determined by simultaneously fitting nanobody binding association kinetic curves obtained at different nanobody concentrations (L) to the equation above with the relationship between *k*_*obs*_ and the two kinetic binding rate constants *k*_*on*_ and *k*_*off*_ given by:

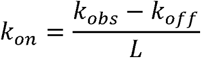

Kinetically determined nanobody K_D_ values were determined with the following equation:

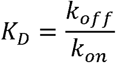

For nanobody dissociation experiments were fitted to the following mono-exponential decay equation:

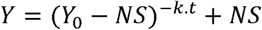

where Y_0_ is the nanobody binding at the time of ligand addition (*t_0_*), NS is the nanobody bound at infinite time, and *k* is the rate constant of nanobody dissociation.

Statistical significance was defined as p<0.05 using an unpaired t-test, or a one-way or two-way ANOVA where appropriate.

