## Supplementary Information and Figures for "Monitoring ligand-induced changes in receptor conformation with NanoBiT conjugated nanobodies"

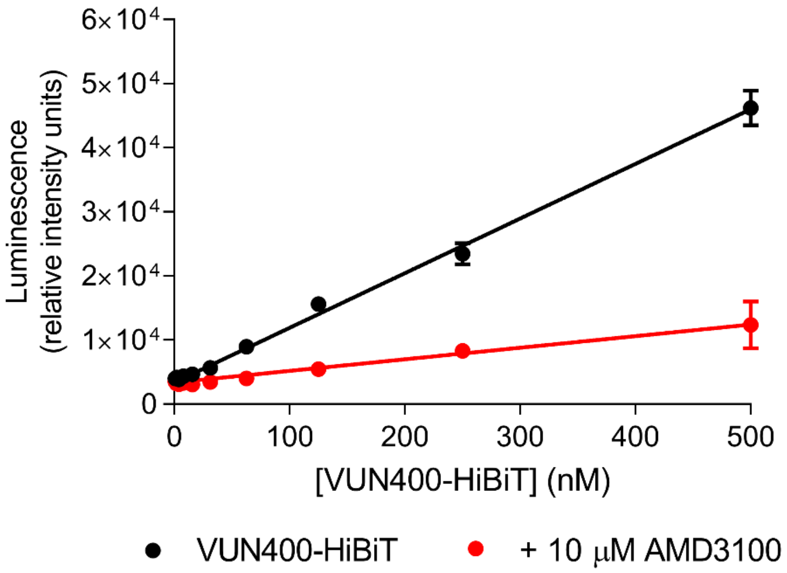


**Supplementary Figure 1 (related to Figure 5).** Saturation binding of VUN400-HiBiT at the SNAP-CXCR4 in the absence (black circles) or presence of 10 µM AMD3100 (red circles). Data are mean ± SEM from triplicate determinations in a single experiment. This single experiment is representative of five separate experiments.

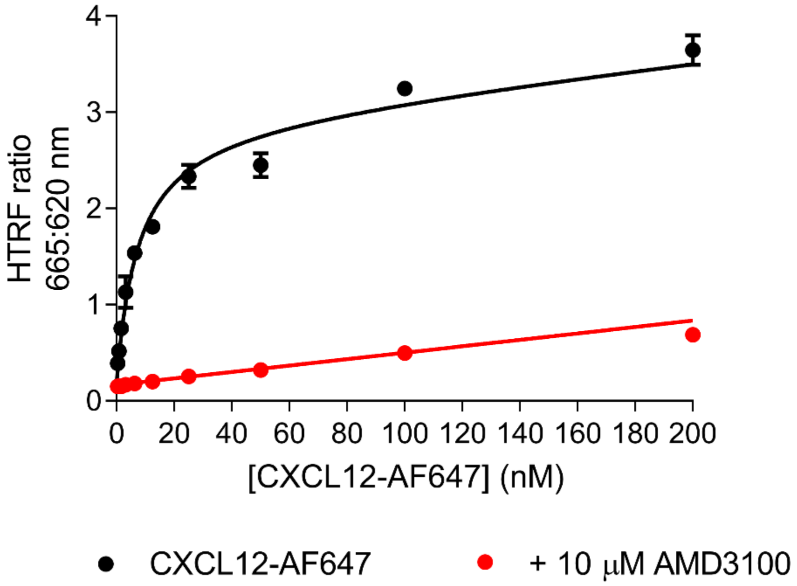


**Supplementary Figure 2 (related to Figure 5).** TR-FRET saturation binding of CXCL12-AF647 at the SNAP-CXCR4 in the absence (black circles) or presence of 10 µM AMD3100 (red circles). Data are mean ± SEM from triplicate determinations in a single experiment. This single experiment is representative of five separate experiments.
